# The mRNA-binding protein HuR is a kinetically-privileged electrophile sensor

**DOI:** 10.1101/2020.04.07.029330

**Authors:** Jesse R. Poganik, Alexandra K. Van Hall-Beauvais, Marcus J. C. Long, Michael T. Disare, Yi Zhao, Yimon Aye

## Abstract

The key mRNA-binding proteins HuR and AUF1 are reported stress sensors in mammals. Intrigued by recent reports of sensitivity of these proteins to the electrophilic lipid prostaglandin A2 and other redox signals, we here examined their sensing abilities to a prototypical redox-linked lipid-derived electrophile, 4-hydroxynonenal (HNE). Leveraging our T-REX electrophile delivery platform, we found that only HuR, and not AUF1, is a kinetically-privileged sensor of HNE in HEK293T cells, and sensing functions through a specific cysteine, C13. Cells depleted of HuR, upon treatment with HNE, manifest unique alterations in cell viability and Nrf2-transcription-factor-driven antioxidant response (AR), which our recent work shows is regulated by HuR at the Nrf2-mRNA level. Mutagenesis studies showed that C13-specific sensing alone is not sufficient to explain HuR-dependent stress responsivities, further highlighting a complex context-dependent layer of Nrf2/AR regulation through HuR.

## Introduction

Regulation of mRNA is key strategy that cells employ to dynamically control gene expression. mRNA-binding proteins (mRBPs) are key players that orchestrate these regulatory programs. Of the hundreds of mRBPs identified in mammalian cells,^[1]^ two of the most crucial mRBPs are HuR and AUF1.^[2]^ These two proteins are widely expressed, and are likely present in the same cells in many instances. These proteins share similar binding preferences (favoring so-called AU-rich sites), and have at least partially-overlapping substrate preferences.^[3]^ Overall these proteins are believed to have opposing effects on their targets: HuR stabilizes its targets whereas AUF1 destabilizes its targets.^[4, 5]^ However, there are several notable exceptions to these observations, and the molecular mechanisms by which each protein functions to regulate specific target mRNAs vary quite considerably. Both proteins are highly relevant to several disease states, including cancers.^[6, 7]^ The mode of action of HuR in cancers is generally tumor promoting, likely through pleiotropic mechanisms, for instance: HuR can promote stability of numerous oncogenic transcripts, such as ERBB-2;^[8]^ HuR can also promote the angiogenic switch;^[9]^ and HuR is also linked to chemoresistance of cancer cells.^[10]^ Unsurprisingly, knockdown of HuR is an established means to reduce anchorage-independent growth and oncogenic potential of at least some sets of cancer cells.^[11, 12]^ The disease relevance of HuR has propelled efforts to identify small-molecule inhibitors of HuR, which have yielded some initial hits that show potential promise as useful therapeutics.^[13, 14]^ These molecules mostly appear to interfere with HuR dimerization, which is essential for HuR function.^[15]^ None of these inhibitors have yet reached even advanced preclinical investigations. Conversely, the role of AUF1 is more nuanced, as it appears to show both tumor-suppressive and tumor-promoting activities, depending on context. Thus, AUF1 inhibitors may also be potentially relevant to therapeutics, although as can be expected, AUF1 targeting is less likely to be a general strategy to combat cancer than HuR. Unsurprisingly, AUF1 inhibitors are far less well developed than HuR inhibitors. Intriguingly in at least one instance, some data indicate that HuR-targeted inhibitors affect AUF1’s mRNA-binding abilities,^[16]^ although this was observed only under certain conditions.

Interestingly, the mRNA-stabilizing and destabilizing abilities of HuR and AUF1 are affected (directly and/or indirectly) by small-molecule stress-inducers ^[17-20]^ For instance, cytoplasmic translocation of HuR, a postulated marker of cellular stress, is promoted by a number of stress-inducing small-molecule stimulants such as H_2_O_2_, arsenite, and the cyclopentenone prostaglandin A2 (PGA2) lipid-derived electrophile housing a Michael acceptor (Figure 1A).^[17, 18]^ Treatment with PGA2 also increases the affinity of HuR for p21 mRNA.^[18]^ Similarly, AUF1 binds and promotes degradation of cyclin D1 and COX-2 mRNA upon PGA2 stimulation.^[19]^ In line with the purported stress-relevance of HuR and AUF1, we recently reported that these proteins modulate the antioxidant response (AR)—a key stress response pathway—by directly regulating the mRNA of Nrf2, the transcription factor responsible for AR upregulation.^[21]^ Nrf2/AR is triggered by some of the same molecules that influence HuR/AUF1 function, particularly electrophiles such as PGA2. However, the link between HuR/AUF1 electrophile regulation and AR also remains untested.

**Figure 1.**
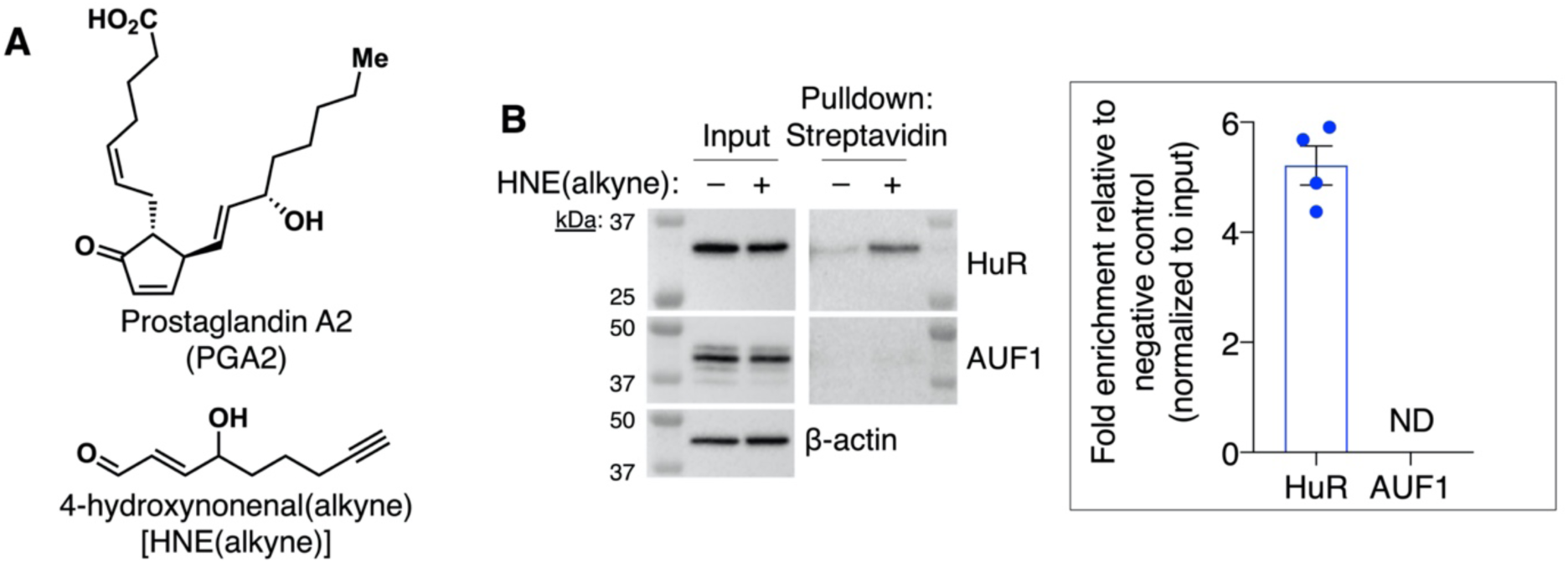
HuR, but not AUF1, is a sensor of HNE in HEK293T cells. (A) Structures of the native RES prostaglandin A2 (PGA2) and alkyne-functionalized HNE. (B) HEK293T cells were treated with HNE(alkyne) (20 µM, 18 h). Click chemistry was used to biotinylate HNE-modified proteins, which were then enriched with streptavidin resin. Shown is a representative blot from 4 independent experiments. Inset at right: Quantification (mean±SEM) of western blot data from n=4 independent replicates. ND, not detected.

The results above raise the possibility that both HuR and AUF1 may contain inherently “ligandable” cysteines that can be leveraged for covalent drug design and to better understand the role of these proteins in electrophile signaling.^[22, 23]^ However, it is important to recognize that the above electrophiles and oxidants are pleiotropic, and hence experimentally-observed outputs can be ascribable to either on-target effects (i.e., direct protein targeting) or off-target effects (i.e. secondary effects that feedback to regulate activity of those proteins), or potentially both. Parsing these different regulation modes is thus particularly challenging. Typically, understanding the role of specific targets in response to electrophiles requires careful and diagnostic experiments performed *in vitro* or under protein-targeted conditions in cells to unravel the site(s) of modification. Specific mutants of the proteins can then be expressed in cells to assign importance of specific on-target protein modification to the response to the specific molecular stressor. However, there has not even been direct evidence furnished regarding the electrophile sensing ability of HuR/AUF1. In the wake of modern-day pharma’s increased interest in design and optimization of covalent drugs, and further with the realization that electrophilic molecules are themselves directly applicable to electrophilic drug design, indications of electrophile sensitivity of these proteins very much warrant further investigation.

In this paper, screening the electrophile-sensing ability of endogenous HuR and AUF1 in controlled manner in HEK293T cells allowed us to establish that only HuR is a sensor of native reactive electrophilic species (RES) represented by the prototypical lipid-derived electrophile, 4-hydroxynonenal (HNE). This result underscores the need to thoroughly address on-target electrophile labeling using unbiased assays. Focusing on HuR electrophile sensing, our T-REX targeted-electrophile delivery platform^[24]^ allowed us to ascribe HuR’s HNE-sensing ability to a single kinetically-privileged cysteine, C13. HuR(C13S) was unable to sense HNE both *in vitro* and in cells. We further found HuR-dependent modulation of Nrf2/antioxidant response (AR)–a key cell defense pathway regulated by HuR^[21]^–upon stimulation of cells with HNE. This effect ultimately transpired to not be fully explained by modification of HuR(C13), although there was a significant difference in response between cells expressing wild type HuR and HuR(C13S). Collectively, these observations add an unexpected layer of complexity to a HuR–AR regulatory program we recently reported,^[21]^ and point to a means to covalently target HuR, orthogonal to AUF1, in future drug discovery endeavors.

## Results

### Initial screen of HNE-sensing ability reveals that HuR, but not AUF1, is an electrophile sensor in cells

We set about investigating the direct RES-sensing capability of HuR and AUF1 in live HEK293T cells. HNE—an endogenous RES with a reactive core chemically similar to that of PGA2 (Figure 1A)—was selected as a representative bioactive RES.^[25, 26]^ We first examined the extent to which endogenous HuR and AUF1 can sense HNE administered to cells globally. Treatment of HEK293T cells with alkyne-functionalized HNE [HNE(alkyne)] (20 μM, 18 h), followed by biotin-azide Click coupling^[27]^ and enrichment of HNEylated proteins using streptavidin pulldown, revealed that HuR, but not AUF1, was sensitive to HNE (Figure 1B). The negative result for AUF1 is contrary to what may have been predicted given the published data, and underscores the need to perform precise identification of electrophile modified proteins.

### HuR senses HNE specifically through C13

HuR is a multidomain protein harboring a total of 3 cysteines (Figure 2A), one of which (C13) resides close to the unstructured N-terminus,^[30]^ and two of which (C245 and C284), also appear to be solvent accessible, at least based on crystal and NMR solution structures of the RRM3 domain wherein these two cysteines reside (Figure 2B).^[31-33]^ Cysteine 13 (C13) of human HuR has been previously proposed to act as a redox sensor that functions through formation of a disulfide bridge between two molecules of HuR. This postulate was based on gel filtration analysis of a truncated HuR construct (Residues 2–189, spanning the unstructured N-terminal region housing C13 and the first two RNA-recognition motif [RRM] domains; Figure 2A). This construct failed to dimerize when C13 was mutated. Although conclusions from these experiments are limited as they study the truncated protein, we first tested the hypothesis that C13 is the electrophile sensor within HuR. The HNE bolus dosing experiments that were used above constitute a potential means to test this hypothesis. However, bolus dosing uses prolonged electrophile exposure, and hence constitutes a rather insensitive means of evaluating electrophile sensitivity in cells, with only the least electrophile sensitive proteins (such as AUF1) likely to show as electrophile insensitive. As our focus now switched to derive more precise information, we chose to use the T-REX assay,^[24]^ a unique means to address this question. T-REX is a particularly stringent test of RES-sensitivity in cells because this assay occurs under RES-limited conditions, mimicking endogenous RES-signaling events in living systems. Furthermore, T-REX directly provides a scalar value that quantifies a specific protein’s electrophile-sensing capacity, called “delivery efficiency”. A variety of observations and theoretical treatments indicate delivery efficiency is independent of protein expression, providing a more robust measure of relative electrophile sensing than most other, if not all, methods available.^[24, 34, 35]^ Thus, delivery efficiency can be compared across different proteins and experiments with more overall confidence than from bolus electrophile dosing experiments. T-REX employs genetic fusion of a HaloTag to the POI.^[36]^ A custom-designed bio-inert precursor to a given RES, in our case, HNE (referred to as “Ht-PreHNE”), is directed specifically to the HaloTag, through a hexylchloride linker which selectively and irreversibly conjugates to HaloTag stoichiometrically *in vivo*. After washing away excess unbound Ht-PreHNE, a brief UV-light illumination (365 nm, 5 mW/cm^2^, 5 min) liberates HNE in the immediate vicinity of the Halo-POI. In the resulting “encounter complex”, competition ensues between inherent reactivity of the POI to HNE and diffusion of HNE from the encounter complex. Provided the POI is a kinetically-privileged sensor of HNE, it will react with the RES before it diffuses away (Figure 2C). The electrophile cannot label the POI after it has diffused away from the complex, even when the POI is particularly electrophile reactive.

**Figure 2.**
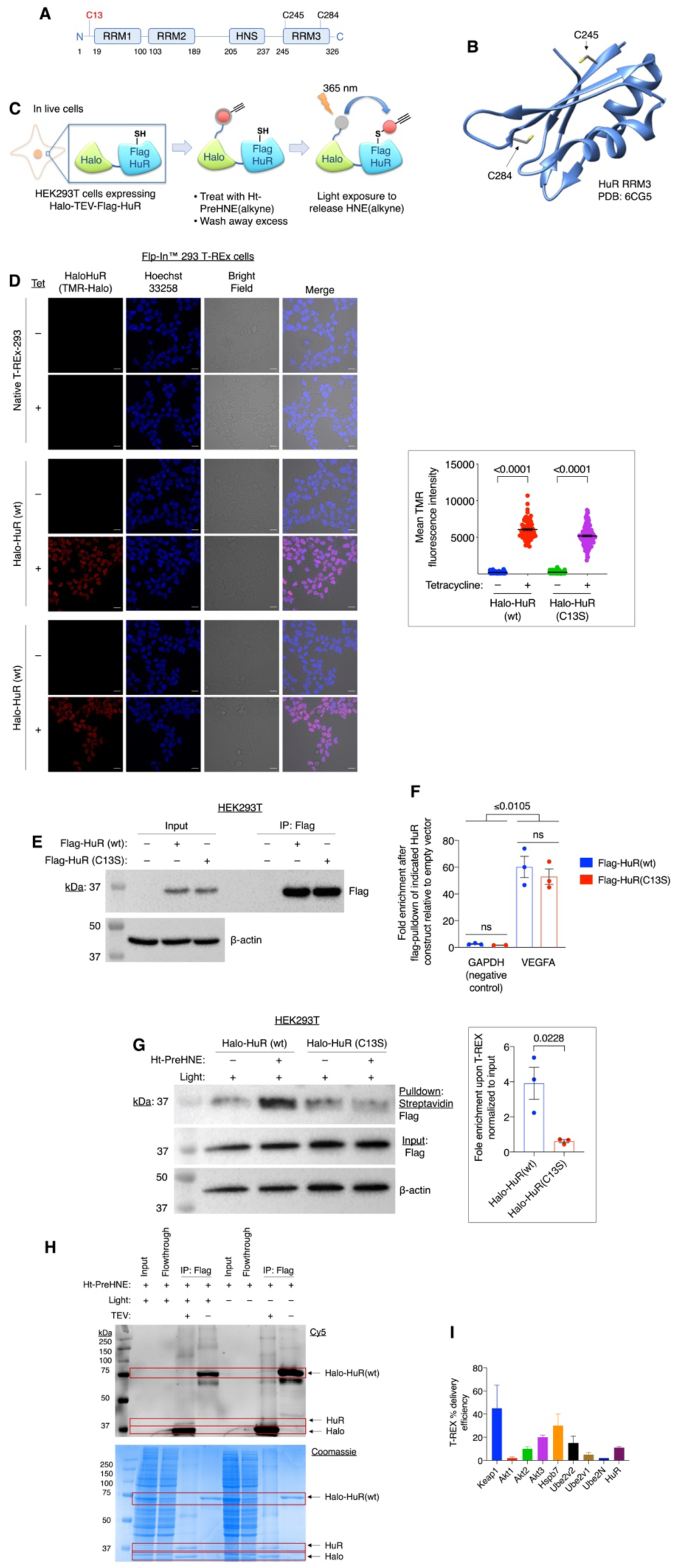
HuR senses HNE specifically through C13; Halo-tagging and mutation of C13 do not affect cellular distribution or mRNA-binding ability of HuR. (A) Domain structure of HuR highlighting its three RNA-recognition motifs (RRM), the HuR nuclear shuttling sequence (HNS) which allows HuR to translocate from nucleus to cytoplasm,^[28]^ and its three cysteine residues. (B) Crystal structure of HuR RRM3, featuring 2 of the 3 cysteines in HuR (PDB: 6CG5). Note that no crystal structure reported to date includes C13, which resides in the unstructured N-terminal region of HuR. (C) Schematic of T-REX HNE delivery platform, which occurs in cells. The protein of interest (POI) is genetically fused to HaloTag and expressed from a plasmid introduced by transient transfection. A bifunctional inert photocaged precursor to HNE [Ht-PreHNE(alkyne)] is introduced and it covalently binds to HaloTag, stoichiometrically. Excess unbound probe is washed away. Then a brief pulse of UV light (365 nm, 5 mW/cm^2^, 5 min) liberates the alkyne-functionalized HNE in proximity to the POI. If the POI were to be a bona fide RES sensor, it would react with the RES in the ensuing “encounter complex” prior to diffusion; otherwise the electrophile would be released to the bulk of the cell, without labeling the target protein. (D) Monoclonal Flp-In T-REx 293 cells with an integrated tetracycline-inducable myc-Halo-TEV-Flag-HuR (“Halo-HuR”, wt or C13S) construct in a defined locus were induced with tetracycline (75 ng/ml) for 48 h and then treated with Halo-TMR (100 nM) overnight. Live cells were counterstained with Hoechst 33258 dye to mark nuclei and imaged using a Zeiss LSM710 confocal microscope. Scale bars, 20 μm. Inset at top right: Quantitation using mean fluorescence of cells determined by the “Measure” tool in ImageJ (mean ± SEM of n>40 cells for each condition). (E) HEK293T cells were transfected with empty vector or Flag-tagged HuR variants as indicated. Flag-HuR was immunoprecipiated and the bound mRNA was isolated. Shown is one representative blot. (F) qPCR analysis of the mRNA recovered from the IP in (E) demonstrates that both Flag-HuR(wt) and Flag-HuR(C13S) bind to VEGFA, a known HuR target,^[29]^ with similar efficiency (mean ± SEM of ≥2 independent replicates). GAPDH serves as a negative, non-target control. (G) Halo-HuR(wt) and Halo-HuR(C13S) mutant were expressed in HEK293T cells. Cells were subjected to T-REX as described in (C), triggering the release of HNE(alkyne) in proximity to HuR. Following HNE delivery, cells were lysed, the Halo domain was cleaved using TEV protease, the lysate was subjected to Click coupling with biotin-azide, and proteins were precipitated and washed thoroughly to remove unbound biotin-azide. Biotinylated proteins were then enriched using streptavidin resin. Halo-HuR(C13S) failed to sense HNE. Inset at right: Quantitation using the “Gel Analysis” tool of ImageJ [n=3 independent replicates (mean ± SEM)]. All p-values were calculated with Student’s t-test. (H) HEK293T cells were transfected with Halo-HuR(wt) for 48 h, then treated with Ht-PreHNE (20 μM; 2 h). Excess Ht-PreHNE was washed away, and cells were exposed to light (365 nm, 5 mW/cm^2^, 5 min), then immediately harvested. Total protein levels were normalized using Bradford assay and Halo-HuR was enriched using Flag IP. Proteins were eluted with Flag peptide, then Click chemistry was used to append Cy5 to modified proteins. Following SDS-PAGE, delivery efficiency was calculated using the following equation:

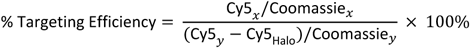

Where Cy5_x_ is the Cy5 signal from modified HuR, Coomassie_x_ is the density of the corresponding Coomassie band, Cy5_y_ is the Cy5 signal from Halo-HuR not exposed to light, Cy5_Halo_ is the Cy5 signal from cleaved Halo exposed to light, and Coomassie_y_ is the density of the Coomassie band for Halo-HuR not exposed to light. Shown is one representative gel from 3 independent experiments; delivery efficiency was determined to be 11 ± 1% (mean ± SEM). (I) Comparison of delivery efficiency for several proteins studied using T-REX.

We first verified that the HaloTag within our Halo-HuR fusion proteins was active by confirming binding to TMR-HaloTag ligand in cells (Figure 2D). Significant TMR signal was only observed in tetracycline-induced cells, and this signal was strongly nuclear-localized, indicating that Halo-tagging did not affect HuR nucleus:cytoplasm distribution (Figure 2D). Both wt and C13S HuR were also expressed at similar levels in cells, based on TMR signal intensity (Figure 2D) and western blot (Figure 2E). Importantly, RNA-binding protein immunoprecipitation (RIP) confirmed that both HuR(wt) and HuR(C13S) bind VEGFA (a known target mRNA of HuR^[37]^; Figure 2E–F) equally well.

Following T-REX in live cells ectopically expressing Halo-HuR, cell lysis, TEV-protease cleavage (that separates HaloTag from HuR), Click coupling with biotin-azide, and streptavidin enrichment, HuR(wt) was enriched 4-fold above HuR(C13S) (Figure 2G). This result demonstrated that C13 is responsible for HNE-sensing. Specifically, the competitive setting deployed in T-REX indicates that C13 is a kinetically-privileged HNE-sensor. To set this sensing in greater context, we calculated the extent of HuR HNE-modification. These experiments showed that the delivery efficiency is 11 ± 1% (Figure 2H). This value is at the lower end of the privileged RES-sensor proteins we have examined under similar conditions in cells, although it is much higher than proteins such as Akt1 and Ube2N, which are not sensors of HNE (Figure 2I).^[24, 34, 38-41]^ The case of HuR is thus interesting as it gives us an opportunity to understand how electrophile signaling occurs on less electrophile-sensitive proteins, perhaps allowing us to test some of our recently-published postulates.^[42]^

We first further validated the privileged RES-sensing property of HuR, and the involvement of C13 in electrophile sensing using purified recombinant HuR. Consistent with numerous reports noting the insolubility and lability of HuR purified from *E. coli*, untagged HuR was poorly soluble and difficult to work with. Noting that our data indicated that Halo-tagged-HuR was active in mammalian cells, we opted to purify Halo-tagged variants of HuR. Consistent with Halo-tag’s reported stabilizing ability,^[43]^ Halo-Tag-HuR and Halo-Tag-HuR(C13S) showed improved stability and solubility (Figure 3A). We next assessed the electrophile sensitivity of the two Halo-Tagged HuR’s, using an assay in which the protein (Halo-Tag-HuR(wt or C13S)) at μM concentrations is mixed with equal amounts of HNE(alkyne). Labeling of the protein is then measured as a function of time, using Click coupling to FAM-azide. Our data revealed a 25-fold difference in time-dependent labeling efficiency between Halo-HuR(wt) and C13S equivalent (Figure 3B). Note: since the Halo protein is present equally in both reaction vessels, and since the starting concentration of HNE was equal in each, this result rules out off-target reaction with Halo as a reason for these results. The outcomes from our T-REX experiments and in vitro HNEylation analysis provide unequivocal evidence that HuR(C13) is a functional HNE-sensor among 3 total Cys residues in human HuR (Figure 2A-B).

**Figure 3.**
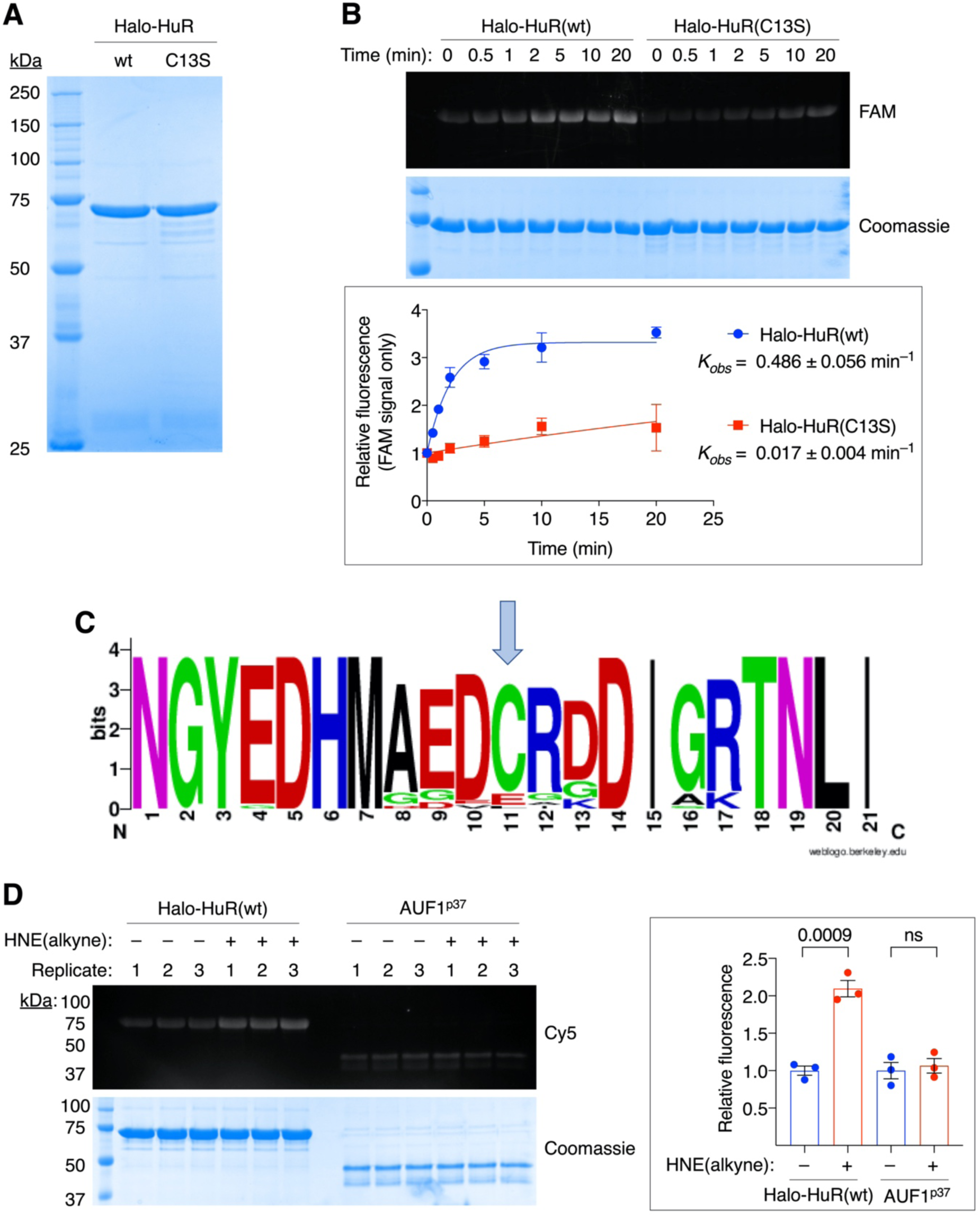
HuR(wt) senses HNE in vitro, whereas HuR(C13S) and AUF1 do not. (A) SDS-PAGE analysis of purified Halo-HuR. Halo-HuR(wt or C13S) was expressed in and purified from *E. coli* using TALON affinity resin followed by size-exclusion chromatography (SEC). 2 μg of each protein was loaded. (B) Recombinant Halo-HuR(wt or C13S) (40 μM) was incubated with equimolar HNE(alkyne) for the indicated time points, upon which a portion of the reaction was diluted 20-fold into chilled buffer. FAM-azide Click followed by SDS-PAGE gel allowed for visualization of modified protein. Inset below: Quantitation using the “Gel Analysis” tool of ImageJ. Fluorescent signal increase (relative to t = 0) was plotted against time and fit to a pseudo-first order binding model [y = 1+2.32(1–e^(–*Kobs*\*x)^)] using Prism. The plateau for both fits was constrained to the saturation plateau for Halo-HuR(wt) (3.32). n=2 independent replicates per point (mean ± SEM). (C) Evolutionary conservation of C13 and flanking residues. The protein sequence of HuR from a selection of about 30 species was aligned to human HuR using MEGA7 and the sequence logo was constructed using WebLogo (UC Berkeley). Arrow indicates the position of C13. (D) Purified Halo-HuR(wt) and AUF1^p37^ (40 μM for each) were incubated with equimolar HNE(alkyne) for 5 min, then immediately diluted 20-fold into chilled buffer. Click chemistry was then used to append Cy5-azide to modified proteins. Coomassie staining was used to visualize protein loading (note Coomassie signal intensities for AUF1 and HuR are not an indication of *molar* amounts of proteins loaded due to the different molar masses of the two proteins; and because Coomassie interacts with different proteins differently). Three independent replicates are shown for each protein. Inset at right: Quantitation of in-gel Cy5 fluorescence using the “Gel Analysis” tool of ImageJ [n=3 independent replicates shown (mean ± SEM)]. All p values were calculated with Student’s t-test.

To further set these data in context, we examined the phylogeny of HuR cysteines compared to other privileged sensors we have investigated previously. HuR(C13) is conserved across higher vertebrates including primates, mice, rats, some birds, and some fish (Figure 3C). However, HuR(C13) is not present in *D. rerio*, a system that we have shown to undergo many electrophile sensing processes that also occur in humans, such as electrophile sensing/signaling by Ube2V2,^[34]^ HSPB7,^[44]^ and Akt3.^[45]^ Similar to the relatively low delivery efficiency, this lack of a clear conservation pattern of HuR(C13) could be taken as evidence that RES-modulation of HuR is a complex mode of regulation that is context dependent.^[46]^ The other two cysteines within HuR are in fact more conserved than C13 giving further credence to this argument.

Finally, we also used this *in vitro* labeling assay to corroborate the lack of HNE-sensing by AUF1 that we observed in cells (Figure 1B). Under conditions in which HNEylation of HuR saturated (5 min; Figure 3B), AUF1 failed to be modified at all (Figure 3D). The consistent outputs between data from T-REX, bolus dosing in cells, and *in vitro* is in overall agreement with a large amount of data previously disclosed from our group.

### HuR knockdown perturbs viability of cells exposed to electrophiles and oxidants, but C13 modification is not fully sufficient to explain this effect

Based on the data above, we hypothesized that depletion of HuR might affect the ability of cells to survive when exposed to redox stressors. We chose to test this hypothesis in cells exposed to H_2_O_2_, a prototypical oxidant, in addition to cells exposed to HNE, which serves as a prototype for electrophilic stress. Growth inhibition experiments carried out in HEK293T cells depleted of HuR versus control cells (treated with untargeted siRNAs) revealed that cells deficient of HuR were 15-20% more sensitive to HNE and H_2_O_2_ than control cells (Figures 4A–B). To set these data more in context, we examined how knockdown of Nrf2 affected the sensitivity of HEK293T cells to the same reagents. We chose Nrf2 as a comparison because this protein is the key antioxidant response transcription factor, which controls a battery of antioxidant defense proteins that are upregulated upon exposure to electrophiles or oxidants.^[47]^ We have also recently shown that expression levels of Nrf2 and response to electrophilic stress are regulated by HuR expression.^[21]^ Knockdown of Nrf2 also rendered cells around 20% more sensitive to HNE (Figure 4C). However, we observed no difference in EC_50_ for growth inhibition by H_2_O_2_ (Figure 4D). Thus, HuR knockdown affects cellular sensitivity to redox stress, at least as significantly as, and in some cases more than, knockdown of Nrf2, a protein that is believed to be essential to protect against various small-molecule stressors. Consistent with this notion, AR levels (which directly and faithfully report on levels of active Nrf2^[21]^) were consistently upregulated by ∼2-fold in cells transfected with siHuR and treated with HNE, compared to siControl-transfected cells (Figure 4E). No modulation of AR was observed on average in cells transfected with siHuR and exposed to H_2_O_2_ (Figure 4F). Importantly, AR levels in non-treated cells were significantly suppressed when HuR was knocked down, validating HuR knockdown (Figure 4G).^[21]^ We note that the antagonistic effects of HuR-knockdown cells under non-treated conditions (AR suppressed) versus HNE-treated conditions (AR hyperstimulated) reflects the pleiotropic nature of both electrophiles and HuR.

**Figure 4.**
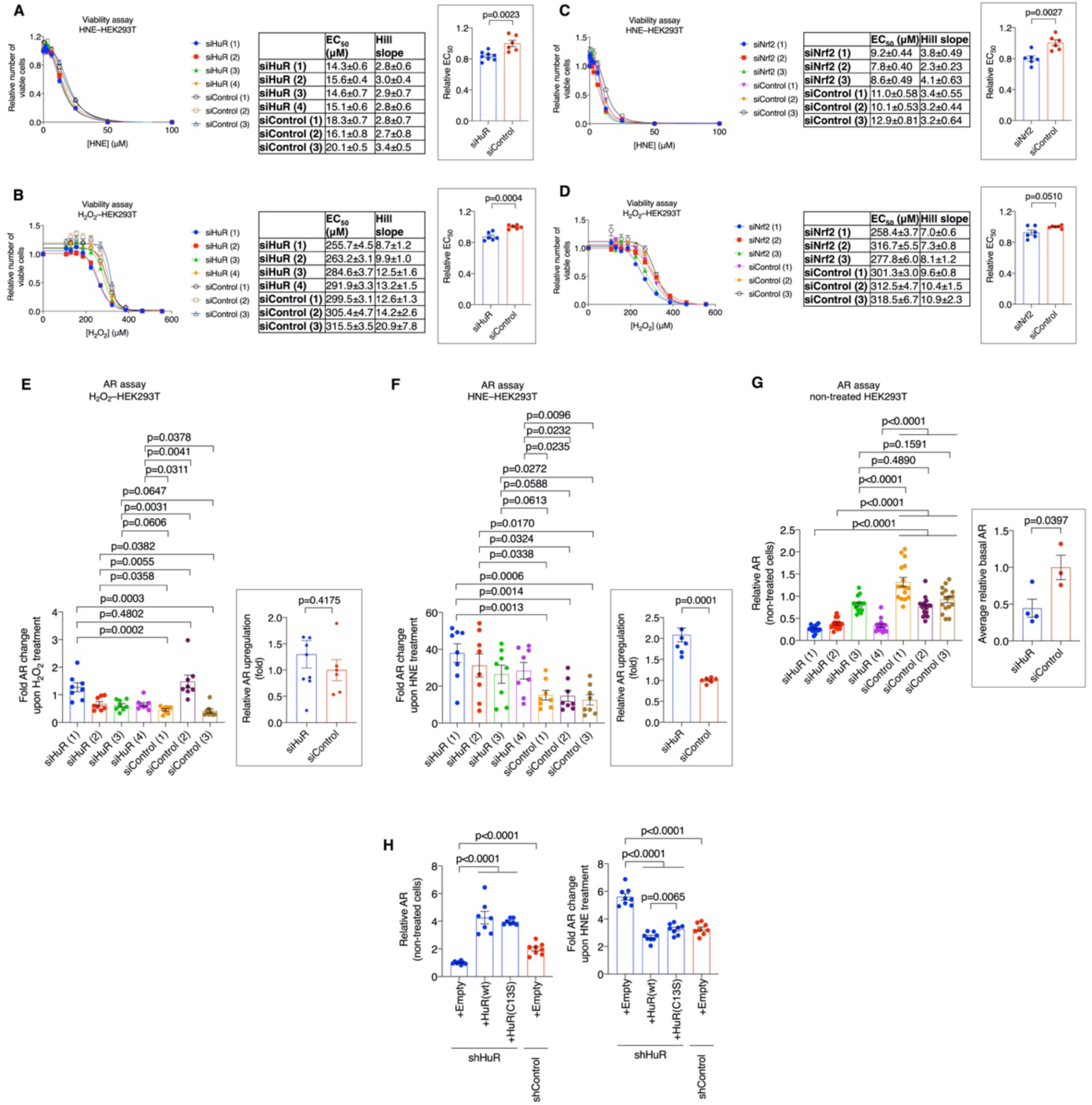
Depletion of HuR sensitizes cells to proliferation inhibition/toxicity induced by HNE; this is likely mediated by HuR regulation of Nrf2. (A–D) AlamarBlue was used to assess growth inhibition of HEK293T cells transfected as indicated and treated for 48h with the indicated molecules (mean±SEM of n=8 for all points except for non-treated, where n=16). All EC_50_ data were fit to the equation *y*=*Bottom*+(*Top*-*Bottom*)/1+(*x*^*Hill slope/*^*EC*_50_^*Hill slope)*^. Table at right in each case shows determined EC_50_s and Hill Slopes: note change in sensitivity is reflected only in EC_50_, not in Hill coefficient. Inset at far right: Global analysis of EC_50_ (mean±SEM of n=8 for siHuR treated with HNE; n=6 for all other conditions). All individual siHuRs and siControls were included in this analysis. (E, F, and G) AR activity of HEK293T cells transfected as indicated and either treated with HNE (E) or H2O^2^ (F), or non-treated (G), respectively (mean±SEM of n=8 per condition). Inset at right: similar analysis of all siHuRs and siControls as in A–D (mean±SEM of n=8 for siHuR and n=6 for siControl). (H) AR activity of shHuR/shControl cells transfected as indicated (mean±SEM of n≥6 per condition). All p values were calculated with Student’s t-test. Note: the inherent batch to batch variations and stabilities of HNE and siRNA transfection render absolute EC_50_’s variable across runs, e.g. A versus B. We thus compare relative effects relative to specific siRNA controls for Nrf2 and HuR.

We then examined the extent to which perturbation of AR in HuR-knockdown cells is dependent on modification of C13. Transfection of HuR(wt) into shHuR-lines fully rescued the suppression of AR observed in unstimulated shHuR-cells transfected with an empty vector relative to identical shControl cells treated with HNE (Figure 4H). This procedure also *reduced* levels of AR upregulation upon HNE-treatment in shHuR-cells to AR-levels observed in shControl cells (Figure 4H, second bar in each set). Transfection of HuR(C13S) was able to fully rescue suppressed AR levels in unstimulated shHuR-cells, again attesting to the activity of HuR(C13S) (Figure 4H, third bar). In these rescued cells, the active HuR species is HuR(C13S), and thus privileged electrophile sensing by HuR cannot occur. We would, therefore, expect these HuR(C13S)-expressing cells to respond to HNE-treatment similarly to HuR-knockdown cells (i.e., show elevated AR in response to HNE), provided elevated AR observed in HuR knockdown cells were due to loss of effects attributable to electrophile labeling at HuR(C13): otherwise we could consider HuR(C13S) cells behaving the same as HuR(wt)-expressing cells, if C13 electrophile labeling were irrelevant to the elevated electrophile response observed in HuR-knockdown cells. Cells where HuR-knockdown was rescued with HuR(C13S) featured *partially* elevated AR levels upon treatment with HNE, compared to cells where HuR-knockdown was rescued with wt HuR. Thus, although HuR regulates AR in response to HNE, modification of C13 is only partly (but *statistically significantly*) able to account for this effect. Furthermore, given that HuR and HNE are pleiotropic drivers of AR, these data do not allow us to assign how HuR(C13)-modification precisely affects HuR activity in an otherwise unperturbed cellular background. Such outcomes are quite common for pleiotropic molecules, and have been reported for molecules as diverse as dimethyl fumarate, benzylisothiocyanante, and HNE. Furthermore, the impact of HuR direct electrophile sensing, in terms of global response, is not dissimilar from the contribution made to signaling outputs by proteins such as Pin1 to the overall electrophile response.^[48]^

## Discussion

We set out this project with two principal goals: (1) to evaluate if AUF1 and/or HuR were sensitive to electrophile labeling; and (2) to understand how any electrophile sensitivity impacted the cellular response to redox stress. We began by exploring the electrophile-sensing abilities of HuR and AUF1. We identified that among the 6 cysteines within these two proteins, HuR(C13) was unique at performing a privileged HNE-sensing function. This result underscores the need to perform careful electrophile sensitivity measurements because although both HuR and AUF1 are influenced by the electrophilic prostaglandin, PGA2,^[17-19]^ neither of these two proteins had previously been shown to be labeled by PGA2 or any other electrophile. Separately, HuR is also responsive to H_2_O_2_ treatment, although these experiments used a truncated protein as the substrate. Critically, the kinetics of labeling by specific reagents and relative sensitivities of these proteins, both of which are critical for assessing physiological relevance, had also been ignored.

We first showed that bolus dosing with HNE labelled HuR but not AUF1. This would typically be taken to imply that HuR is a good sensor of HNE. However, such results overall ignore kinetics of protein labeling, offering little insight into how reactive HuR really is. We resolved these questions using T-REX and *in vitro* experiments. Intriguingly, T-REX delivery did not achieve particularly high HNE occupancy on HuR. The 11 ± 1% modification efficiency of HuR achieved under T-REX is not as high as we have observed for several other RES-sensor proteins under T-REX including Keap1 (20–55%, with Halo-fusion at either N- or C-terminus), ^[24, 38-40]^ HSPB7 (30 ± 10%),^[49]^ Akt3 (20 ± 5%),^[41]^and Ube2V2 (15± 6%)^[34]^ (Figure 2I). Consistent with these observations, labeling of HuR (40 μM) by HNE (40 μM) *in vitro* took longer time to saturate than other proteins we have reported to sense HNE by T-REX [such as labeling of Ube2V2 by HNE^[34]^ (approximately 1 min, concentrations of both Ube2V2 and HNE = 12 μM); and HSPB7^[44]^ (2-4 min, concentrations of both HSPB7 and HNE = 12 μM)]. This longer time to saturation occurred despite the fact that concentrations of HuR and HNE were higher than we have typically used (assuming the reaction is second order, *t*_1/2_ is inversely proportional to the concentration of HuR/HNE). Nevertheless, the rate of HuR labeling *in vitro* remains faster than what we have observed for previously reported electrophile-sensitive proteins, and proteins that were not electrophile sensitive under T-REX (e.g. Ube2N). This outcome overall gives us more evidence that delivery efficiency is a useful variable that can be compared between runs and inform on intrinsic electrophile sensitivity in an unbiased manner. Further investigations should shed more light into the overall usefulness of this parameter and how delivery efficiency compares to *in vitro* electrophile sensitivity.

Regardless of the relative electrophile sensitivity of HuR compared to other highly reactive electrophile sensors, it is important to note that we have identified a covalent site on HuR that is orthogonal to AUF1. As we mentioned in the introduction and have written about at length elsewhere,^[45, 50]^ these sites, particularly when embedded in one protein within a group of proteins with overlapping functions, such as HuR and AUF1, are *potential* gold mines for covalent drug design. With covalent inhibitors being on a continued upward trajectory in terms of their importance in drug design, and with HuR being a known anti-cancer target with no approved drugs, this discovery is a first step towards generating new selective HuR inhibitors. This function can be harnessed independent of electrophile signaling functions of C13 that we investigated later.

We proceeded to evaluate how HuR(C13) affected AR response upon electrophile exposure. Our data, which used relatively low concentrations of HNE, for 18 h, showed overall a modest importance of HuR(C13) in the regulation of AR by HuR. Clearly this result shows that the elevated AR upregulation to HNE treatment seen in the HuR-knockdown backgrounds are only marginally attributable to loss of HuR-electrophile sensing. Such outcomes are not unexpected for pleiotropic electrophiles. Although not optimal, it is critical to understand when interpreting these data that, aside from regulating many different proteins, electrophile labeling can impact any function of the POI, including potentially upregulating activities that are not present in the unmodified state. Thus, electrophile signaling is particularly complex for multifunctional proteins, like HuR. Thus, the importance of HuR(C13)-electrophile labeling may only become clear as the regulatory roles of HuR become further elucidated. Critically, we have previously shown that AR regulation by HuR occurs through several functions,^[21]^ perhaps intrinsically mitigating how HuR electrophile signaling can impact AR. It is of course possible that some subset of the HuR interactome could be regulated more acutely by HuR-specific electrophile signaling. It is also important to note that in a number of disease states such as Alzheimer’s and Parkinson’s disease, HNE production is chronically upregulated; HNE adducts have been detected from patient tissue samples.^[51]^

Under these conditions of chronic HNE exposure, which are difficult to model in cultured cells, it is possible that the stoichiometry of RES:HuR covalent modification may increase, potentially giving rise to HuR electrophile labeling becoming more important in regulating AR-signaling behavior. These postulates can all be investigated in detail at a later stage, in further work, although they are outside the scope of this paper. Nevertheless, we were able to ascribe a minor component of HuR’s regulation of AR to modification at C13, which as we discussed above is the typical output, even when studying proteins with a single ascribed activity.

## Experimental Section

### Statistics and data presentation

For experiments involving cultured cells, samples generated from individual wells or plates were considered biological replicates. In the figure legend for each experiment, the number of independent biological replicates and how the data are presented in the figure (typically mean ± SEM) are clearly indicated. P-values calculated with Student’s two tailed unpaired t-test are clearly indicated within data figures. Data were plotted and statistics generated using GraphPad Prism 7 or 8.

### Reagents

HNE(alkyne)^[49]^ and Ht-PreHNE(alkyne)^[24]^ were synthesized as previously described. Unless otherwise indicated, all other chemical reagents were bought from Sigma at the highest availability purity. TCEP was from ChemImpex. Hygromycin was from Invitrogen. AlamarBlue was from Invitrogen, and was used according to the manufacturer’s instructions. Minimal Essential Media, RPMI, Opti-MEM, Dulbecco’s PBS, 100X pyruvate (100 mM), 100X nonessential amino acids (11140-050) and 100X penicillin streptomycin (15140-122) were from Gibco. Protease inhibitor cocktail complete EDTA-free was from Roche. TALON (635503) resin was from Clontech. Ni-NTA resin was from Qiagen. 2020 and LT1 transfection reagents were from Mirus. PEI was from Polysciences. Hoechst dye was from Invitrogen. TMR-Halo ligand was from Promega. Venor GeM PCR-based mycoplasma detection kit was used as stated in the manual and was from Sigma. ECL substrate and ECL-Plus substrate were from Pierce and were used as directed. Acrylamide, ammonium persulfate, TMEDA, Precision Plus protein standard were from Bio-Rad. All lysates were quantified using the Bio-Rad Protein Assay (Bio-Rad) relative to BSA as a standard (Bio-Rad). PCR was carried out using Phusion Hot start II (Thermo Scientific) as per the manufacturer’s protocol. All plasmid inserts were validated by sequencing at Cornell Biotechnology sequencing core facility or Microsynth. All sterile cell culture plasticware was from CellTreat. Antibodies were previously validated^[21]^ using western blot/immunofluorescence to show that multiple independent shRNAs/MOs significantly reduced the levels of the detected protein.

### Cell culture

HEK293T (obtained from ATCC) and Flp-In-293 (Invitrogen) were cultured in MEM (Gibco 51090036) supplemented with 10% v/v fetal bovine serum (Sigma), penicillin/streptomycin (Gibco), sodium pyruvate (Gibco), and non-essential amino acids (Gibco) at 37°C in a humidified atmosphere of 5% CO2. Medium was changed every 2-3 days. HEK293T cells expressing shHuRs/shControls were generated as previously described.^[21]^

### Cloning

Mammalian and bacterial expression plasmids were cloned as previously described, using the Hawaii method.^[21]^ Briefly, the gene of interest was amplified using the forward and reverse primers listed below, then extended and inserted into pCS2+8 or pET28a linearized with EcoRI (NEB). All inserts were fully sequenced by Sanger sequencing at the Cornell Genomics Core Facility.

Site-directed mutagenesis was carried out by PCR amplification of the starting plasmid with forward and reverse mutagenesis primers containing the desired mutation (Table 1) followed by DpnI (NEB) treatment. shRNA-resistant expression plasmids (for rescue experiments) were also produced using this procedure. These code for the same protein but contain mismatches (highlighted in red in the primer sequences; Table 1) in the shRNA-targeting sequence allowing expression of the ectopic protein in knockdown cells. Successful mutagenesis was confirmed by Sanger sequencing.

**Table 1:**
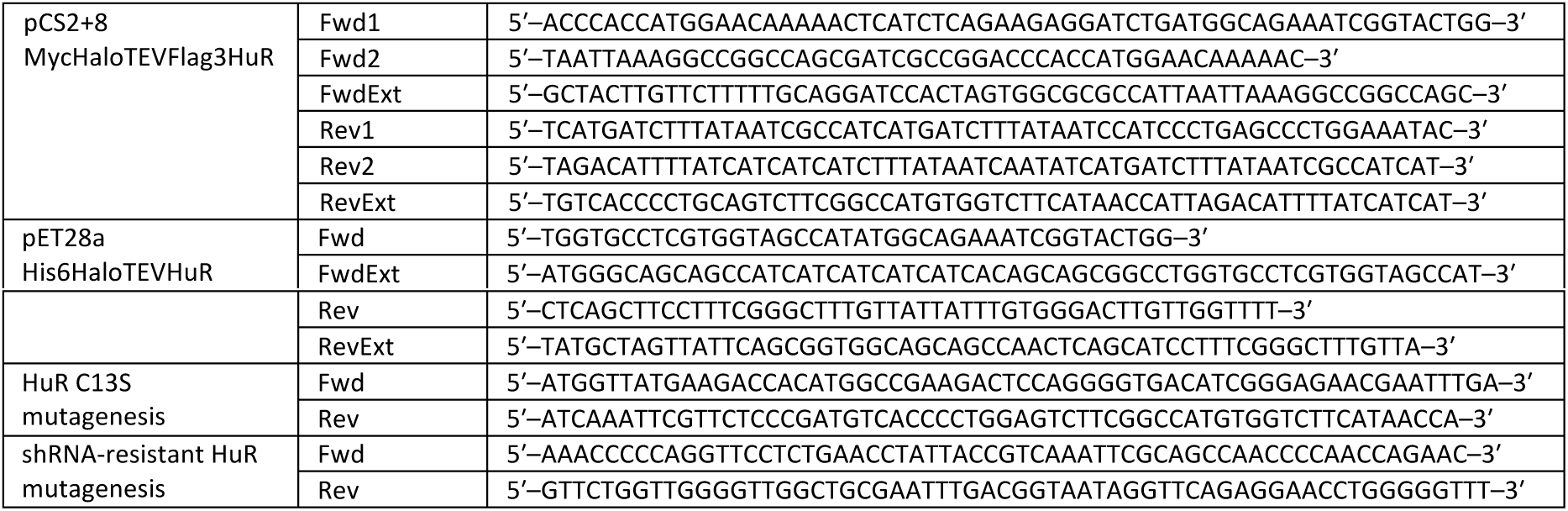
Sequences of cloning and mutagenesis primers.

PEI transfection of HEK293T cellsHEK293T cells (4.5 × 106) in 10 cm diameter plates were transfected with 8 μg total plasmid DNA per plate with PEI (21 g per plate). Media was changed 24 h-post transfection and the cells were incubated 48 h total. For smaller plate sizes, the volumes were scaled accordingly.

### Knockdown of HuR with siRNA

SiRNAs targeting human HuR were obtained from Dharmacon and non-targeting control siRNAs (Control siRNA-A, C, and E) were obtained from Santa Cruz Biotechnology. HEK293T (3.6 × 105 cells) in 6-well plates were transfected with siRNA using Dharmafect I (Dharmacon) for 48 h following the manufacturer’s protocol, then assayed. For co-transfection of siRNA and plasmid(s) for reporter assays, cells were transfected with Dharmafect Duo (Dharmacon) for 48 h following the manufacturer’s protocol, then assayed.

**Table 2:**
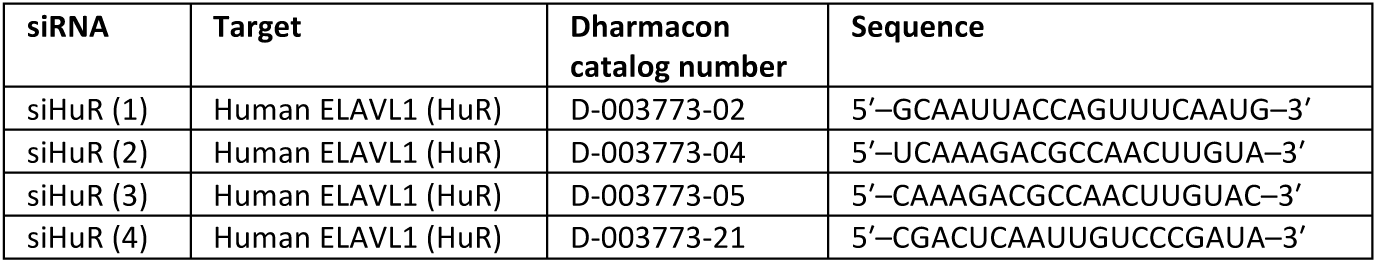
Sequences of siRNAs targeting HuR.

### Western blotting

Cells were resuspended in lysis buffer (50 mM HEPES pH 7.6, 1% Triton X100, 0.3 mM TCEP, 1X Roche cOmplete tablet) and lysed by three cycles of rapid freeze-thaw. Lysates were cleared by centrifugation (20000 × g, 10 min, 4°C) and total protein concentration was determined by Bradford assay (against BSA standard). 50 μg of total protein was loaded per lane, separated by SDS-PAGE, transferred to PVDF, then the membrane was blocked, and incubated with the appropriate antibodies. Detection was carried out on a ChemiDoc-MP imaging system (BioRad) or a Fusion FX imager (Vilber) using ECL Western Blotting Substrate (Pierce) or SuperSignal West Femto Maximum Sensitivity Substrate (Thermo Scientific). Western blot data were quantitated using the Gel Analysis tool in ImageJ (NIH). Bands of interest were integrated and normalized to the loading control.

**Table 3:** Antibodies.

| Antibody | Source | Dilution(s) |
| --- | --- | --- |
| Mouse monoclonal anti-HuR | Santa Cruz Biotechnology sc-5261 | 1:1000 |
| Rabbit polyclonal anti-AUF1 | EMD Millipore 07-260 | 1:500 |
| Mouse monoclonal anti-FLAG M2 | Sigma F1804 | 1:2000 |
| Mouse monoclonal anti-β-actin (peroxidase-conjugated) | Sigma A3854 | 1:40000 |
| Goat polyclonal anti-rabbit IgG (HRP-conjugated) | Cell Signaling Technology #7074 | 1:3000 (WB secondary) |
| Goat polyclonal anti-mouse IgG (HRP-conjugated) | Abcam ab6789 | 1:8000 (WB secondary) |
| Anti-mouse IgG VeriBlot for IP secondary antibody (HRP-conjugated) | Abcam ab131368 | 1:1000 (IP WB secondary) |

aWB, western blot; IP, immunoprecipitation

### Streptavidin pulldown of HNEylated proteins following bulk HNE(alkyne) exposure

HEK293T cells were grown to ∼70% confluence and treated with HNE(alkyne) (20 μM) for 18 h. For experiments involving ectopic expression, cells were transfected using the PEI protocol described above. At 30 h post-transfection, cells were treated with HNE(alkyne) (20 μM) and incubated for a further 18 h.

Cells were then harvested by trypsinization, washed twice with PBS and once with 50 mM HEPES pH 7.6, and resuspended in lysis buffer (50 mM HEPES pH 7.6, 1% Nonidet-P40, 0.2 mM TCEP (Gold Biotechnology), and Roche cOmplete EDTA-free Protease Inhibitor Cocktail). A small amount of glass beads (Sigma) were added and cells were lysed by three rapid freeze-thaw-vortex cycles. Debris was removed by centrifugation (20000 x g, 10 min, 4°C) and protein levels were normalized using Bradford assay. 4-5 mg of total protein was diluted to 0.75 mg/ml in lysis buffer and Click chemistry was used to couple HNE(alkyne)-modified proteins with biotin-azide. The Click reaction contained (final concentrations) 1% SDS (VWR), 5% t-butanol (Fisher), 200 μM biotin-azide (Quanta), 2 mM TCEP (Gold Biotechnology), 1 mM CuSO_4_ (Fisher), 0.1 mM Cu(II)-TBTA (Lumiprobe) and was incubated for 40 min at 37°C. EtOH was added to the reaction mixture to a final concentration of 80% and proteins were precipitated overnight at –80°C. Protein was pelleted by centrifugation (20000 x g, 1.5 h, 4°C), and the pellet was washed as follows: 1 wash with 70% EtOH, 2 washes with 100% EtOH, 1 wash with 100% acetone. The pellet was dried briefly and then resuspended in 8% LDS (Chem-Impex) in 50 mM HEPES pH 7.6 with vortexing and sonication. The resuspended mixture was centrifuged (20000 x g, 10 min, room temperature) to remove insoluble material, and then diluted in 50 mM HEPES pH 7.6 to a final LDS concentration of 0.5%. 50 μL of high-capacity streptavidin agarose (Pierce) was added and the tubes were rotated end-over-end for 3 h at room temperature. The resin was washed three times with 0.5% LDS in 50 mM HEPES pH 7.6 (30 min each with rotation), and proteins were eluted in SDS loading buffer for 15 min at 50°C. Modified proteins were detected by western blotting using antibodies to the protein of interest.

### Streptavidin pulldown of HNEylated proteins following T-REX delivery of HNE(alkyne)

This procedure was adapted from a previous report ^[41]^. HEK293T were transfected with Halo-HuR constructs for 48 h using the PEI transfection protocol above. All further steps were carried out under red-filtered light. Cells were incubated with Ht-PreHNE(alkyne) (15 μM) for 2 h in serum-free media. Cells were then rinsed with serum free media (3 rinses, 30 min each) and exposed to light (365 nm; 5 mW/cm2) for 5 min. Cells were immediately harvested and washed twice with PBS and once with 50 mM HEPES pH 7.6. Cells were resuspended in lysis buffer (50 mM HEPES pH 7.6, 1% Nonidet-P40, 0.2 mM TCEP, and Roche cOmplete EDTA-free Protease Inhibitor Cocktail) and lysed by three rapid freeze-thaw cycles and debris was cleared by centrifugation (20000 x g, 10 min, 4°C). Protein levels were measured using Bradford assay. 0.5 mg of total protein at a concentration of 2 mg/ml was treated with TEV protease (0.3 mg/ml final concentration) for 20 min at 37°C. The protein was then diluted to 0.75 mg/ml and biotin-azide was appended to modified proteins using Click chemistry as described above. All further pulldown steps were performed as described above.

### RIP-PCR

RIP-PCR was carried out following previously described methods ^[52]^. HEK293T cells (4.5 × 106) in 10 cm diameter plates were transfected with the indicated constructs (mixed 1:3 with empty plasmid) or empty plasmid alone (8 μg total DNA per plate) with PEI (21 g per plate). Media was changed 24 h-post transfection and the cells were incubated 48 h total. Cells were washed once with PBS (Invitrogen) and harvested by trypsinization, and washed thoroughly with PBS.

Cell pellets were resuspended in polysome lysis buffer [10 mM HEPES (Chem-Impex) pH 7.0, 100 mM KCl (Fisher), 5 mM MgCl_2_ (Fisher), 0.5% Nonidet-P40, cOmplete EDTA-free Protease Inhibitor Cocktail (Roche), 0.2% vanadyl ribonuceloside complex (NEB)] and frozen at –80°C for at least 30 min. The lysate was thawed, centrifuged twice (20000 × g, 10 min each), and the protein concentration was determined using Bradford Assay. 3 mg of total protein was diluted to 0.5 mg/mL in NT2 buffer [50 mM Tris pH 7.4, 150 mM NaCl (Fisher), 1 mM MgCl_2_ (Fisher), 0.05% Nonidet-P40, 0.2% vanadyl ribonuceloside complex] and incubated with 50 μL of Flag M2 agarose (Sigma) for 3 h at 4°C with end-over-end rotation. The resin was washed at least 4 times (5 min per wash) with NT2 buffer containing 300 mM NaCl and a portion of the resin was retained for western blot analysis. The resin was resuspended in 100 μL of NT2, supplemented with 0.1% SDS and 3 mg/mL proteinase K (Santa Cruz Biotechnology), and incubated at 55°C for 30 min. RNA was isolated using Trizol reagent following the manufacturer’s protocol and analyzed by qRT-PCR as described below.

### Quantitative real time PCR (qRT-PCR)

This was carried out as previously described ^[21, 39]^. Briefly, cells were harvested by aspiration of growth medium and direct addition of Trizol reagent into culture plates. Total RNA was then isolated following the manufacturer’s protocol. 1 μg of total RNA (purity/integrity assessed by agarose gel electrophoresis and concentration determined by A260nm using a BioTek Cytation3 microplate reader with a Take3 accessory) was treated with amplification-grade DNase I (Invitrogen) and reverse transcribed using Oligo(dT)20 as a primer and Superscript III Reverse Transcriptase (Life Technologies) following the manufacturer’s protocol. PCR was performed for two technical replicates per sample using iQ SYBR Green Supermix (Bio-Rad) and primers specific to the gene of interest following the manufacturer’s protocol. Amplicons were chosen that were 150–200 bp in length and had no predicted off-target binding predicted by NCBI Primer BLAST. For genes with multiple splice variants, primers were chosen that amplified conserved sequences across known splice variants. Primers were validated using standard curves generated by amplification of serially-diluted cDNA; primers with a standard curve slope between –0.8 and 1.2 and R2≥0.97 were considered efficient. Single PCR products were confirmed by melting analysis following the PCR protocol. Data were collected using a LightCycler 480 (Roche). Threshold cycles were determined using the LightCycler 480 software. Samples with a threshold cycle >35 or without a single, correct melting point were not included in data analysis. Normalization was carried out using a single housekeeping gene as indicated in each dataset and the ΔΔCt method.

### Expression and purification of proteins

Halo-HuR(WT or C13S) and AUF1p37 were expressed and purified as previously described.^[21]^ TEV protease was expressed and purified as previously described.^[24]^ Concentration of proteins was determined by measuring absorbance at 280 nm and calculating with the extinction coefficient determined by the ProtParam tool of ExPasy. Concentrations determined by this method were also independently confirmed using Bradford assay (BSA standard).

### In vitro HNEylation

All steps were carried out under red-filtered light. Recombinant WT or C13S Halo-HuR (40 μM) or AUF1p37 was treated with equimolar HNE(alkyne) at 37°C for the indicated time. At each time point, a portion of the reaction mixture was diluted 20-fold into ice-cold 50 mM HEPES pH 7.6 with 1% Triton X-100. After the final time point, Click chemistry was used to append FAM-azide or Cy5-azide (as indicated in figures) (Lumiprobe) to modified proteins, and the extent of modification was assessed using in-gel fluorescence. Gels were quantitated using the “Gel Analysis” tool of ImageJ and the data were fit to Equation 2 to derive the time-dependent HNE-modification rate using Prism 7.

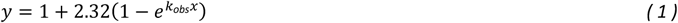

### Analysis of T-REX modification efficiency for HuR

HEK293T cells (4.5 × 106) in 10 cm diameter plates were transfected with Halo-HuR(wt) using PEI as described above. All further steps were carried out under red-filtered light. Following T-REX delivery of HNE as described above, cells were lysed, debris was cleared by centrifugation (20000 x g, 10 min, 4°C), and protein levels were normalized using Bradford assay. Halo-HuR was then enriched by incubation with anti-Flag M2 agarose for 1 h at room temperature. Following elution for 1.5 h with 0.3 mg/ml 3xFlag peptide (ApexBio), eluted proteins were cleaved with TEV protease (0.25 mg/ml final concentration) and subjected to Click coupling with Cy5-azide as described above. The mixture was then run on 15% SDS-PAGE to resolve HuR from Halo. In-gel fluorescence intensity signal (collected with a BioRad Chemidoc MP) was quantitated using the Gel Analysis tool of ImageJ and T-REX targeting efficiency was calculated as previously described ^[24]^ using the equation:

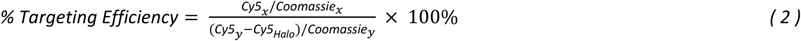

### Growth inhibition assays

HEK293T cells (3000 cells per well) were seeded in 96-well plates. After 24 h, cells were treated with the indicated molecules at the indicated concentrations and incubated for 48 h. AlamarBlue was added to each well and the cells were incubated for a further 4 h, after which fluorescence (excitation 560 nm; emission 590 nm) was measured using a BioTek Cytation 3 microplate reader.

### Luciferase reporter assays for measurement of antioxidant response (AR)

These assays were carried out as previously described^[24]^. Briefly, cells were co-transfected with a 1:0.025:1:1 mixture of pGL4.37 E364A [(ARE:firefly luciferase) Promega]: pGL4.75, E693A [(CMV:Renilla luciferase) Promega]: pCS2+8 Halo-TEV-Keap1: pcDNA3 myc-Nrf2 with Mirus 2020 following the manufacturer’s protocol for 24 h. Cells were then subjected to T-REX (incubated with Ht-PreHNE(alkyne) (15 μM) for 2 h in serum-free media. Cells were then rinsed with serum free media (3 rinses, 30 min each) and exposed to light (365 nm; 5 mW/cm2) for 5 min) and incubated for a further 18 h. Media were removed and cells were lysed for 15 min in passive lysis buffer (Promega) with gentle shaking and homogenized lysate was transferred to the wells of an opaque white 96-well plate (Corning). Firefly and Renilla luciferase activity were measured sequentially on a BioTek Cytation3 microplate reader.

## Acknowledgements

American Heart Association Predoctoral Fellowship (17PRE33670935 to JRP); Swiss National Science Funding (SNSF) Project Funding and SPARK (310030_184729; CRSK-3_190192)); NCCR Chemical Biology (SNSF); NIH Director’s New Innovator (1DP2GM114850); Novartis Medical-Biological Research Foundation (Switzerland); Swiss Federal Institute of Technology Lausanne (EPFL to YA). Mr. Paul Huang and Ms. Yiran Wang are acknowledged for their assistance in photocaged probe synthesis.

## Author Contribution Statement

J.R.P., M.J.C.L., and Y.A. designed the experiments. J.R.P. performed all cell-based assays. A.K.V.H.B., M.T.D., and J.R.P. collaborated on experiments involving gene cloning, protein isolation, and experiments involving purified proteins. Y.Z. performed chemical synthesis. All authors analyzed corresponding experimental data. Y.A. oversaw the project supervision and funding acquisition. J.R.P., M.J.C.L., and Y.A. wrote the manuscript and all authors assisted with proof-editing.

### Entry for the Table of Contents

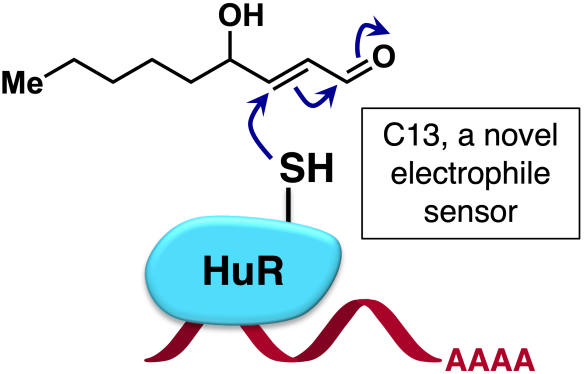

